# Filling the Void: Rapid Revascularization via Vasculogenic Assembly in Semi-synthetic Granular Hydrogel Grafts

**DOI:** 10.64898/2026.06.15.732497

**Authors:** Michael M. Hu, Despina I. Pavlidis, Courtney Lestock, Gonzalo Anyosa-Galvez, Kaley Lollis, Yimeng Zhao, Firaol S. Midekssa, Robert N. Kent, Ariella Shikanov, Brendon M. Baker

## Abstract

Rapid revascularization is critical to tissue graft survival, as delayed reperfusion drives tissue ischemia and compromises cell viability and graft function. Although bulk hydrogels have been explored for promoting vessel formation, vascularization remains too slow to prevent ischemic injury to grafted tissues, highlighting the need for biomaterial platforms that accelerate graft revascularization and reperfusion. In this study, we present granular hydrogel composites (GHCs), where interstitial space is filled with fibrin and collagen to provide a vasculogenic matrix environment. GHCs supported the assembly of embedded endothelial cells into interconnected, lumenized networks in vitro which anastomosed with host vasculature and were systemically perfused 7 days after implantation. Careful optimization studies revealed that GHCs formed from covalently interlinked, RGD-functionalized microgels of 115 µm diameter best supported vascular network formation *in vitro* and intravascular blood perfusion *in vivo*. To test the utility of GHCs for the vascular integration of a demanding and therapeutically relevant parenchymal tissue, GHC-based ovarian tissue grafts were implanted in a murine xenograft model and successfully connected to host vasculature, restoring blood flow to embedded human ovarian tissues within 10 days post-implantation. Notably, endothelial cells seeded within GHCs formed viable vasculature without pre-culture. This work establishes GHCs as a biomaterial platform to rapidly connect parenchymal tissues to host vasculature, with broad translational potential across engineered tissue grafting applications.

## INTRODUCTION

Rapid and sufficient vascularization remains a critical challenge for the survival and efficacy of engineered tissue grafts. While advances in engraftment have improved outcomes for relatively thin tissues where biotransport via diffusion is sufficient (e.g. cartilage, cornea, or skin), transplantation of thicker, more metabolically demanding tissues (e.g. cardiac, hepatic, pancreatic, or ovarian grafts) face high rates of attrition rates due to insufficient nutrient and oxygen transport post-implantation. Inadequate biotransport drives tissue ischemia and the loss of key cell populations, thereby lowering graft efficacy and longevity [1–6]. While successful approaches exist that promote host vasculature angiogenic ingrowth into transplanted tissue grafts, this process typically takes at least 3 weeks, during which parenchymal cell survival and graft function are compromised by unmet metabolic demands.

Graft pre-vascularization, the strategy of incorporating vascular networks prior to implantation, has been shown to improve tissue graft integration with host vasculature, functionality, and longevity [7–9]. Prior work establishing the promise of this approach predominantly used soft and rapidly degradable natural hydrogels (e.g. low-density collagen or fibrin) as scaffolds, due to the well-established ability of these materials to support vascular network assembly. However, these natural biopolymer-based scaffolds tend to be mechanically weak and rapidly resorb following implantation, complicating surgical handling and suitability for applications requiring long-term graft function such as in organ replacement therapies [10–12]. In contrast, synthetic hydrogels are chemically defined, exhibit tunable mechanical properties and degradation characteristics, and are easily modified to provide critical biochemical and biophysical cues required for proper parenchymal cell function. However, typical bulk or monolithic hydrogels formed by crosslinking synthetic polymer backbones yield nanoporous ultrastructures with pore diameters well below 100 nm [13,14]. Given that cells are far larger, such synthetic bulk hydrogels hinder cell spreading and migration, as well as nutrient diffusion [15]. For vasculogenic assembly, a commonly employed strategy for graft pre-vascularization at the capillary scale, endothelial cells must spread and directionally extend protrusions to form intercellular adhesions [16]; as a result, the nanoporosity of bulk synthetic hydrogels slows or in some cases completely prevent microvascular network assembly, thus limiting their utility for pre-vascularized tissue grafts.

Over the past decade, granular hydrogels have been established as a highly modular approach to creating hydrogel-based scaffolds with interconnected microporosity, in contrast to traditional bulk hydrogels [17–19]. Composed of small granular microscale hydrogels (here termed microgels) packed and held together via physical jamming or various chemical crosslinking strategies, these scaffolds intrinsically possess interconnected, microscale porosity [20,21]. This pore architecture not only permits cell invasion into the graft without requiring degradation of the microgels but also provides critical space for the potential formation of vascular networks within the graft. Prior work has shown that ECs can be cultured within granular hydrogels, but these studies have largely yielded aberrant or nascent vascular morphologies rather than functional tubular assemblies [22–26]. Given that endothelial cell morphology is critical to vessel function, here we establish granular hydrogel composites (GHCs) - MAP hydrogels where the pore space is filled with a fibrin-collagen hybrid hydrogel - as a platform that supports rapid EC assembly into a perfusable, tubular vascular network forming simultaneously throughout the graft. We also examined interstitial space dimensions in GHCs by tuning microgel diameter and found larger interstitial geometry allowed for faster vascular network assembly and perfusability. Finally, we investigated whether this approach would enhance vascular integration of ovarian tissues grafts for the treatment of primary ovarian insufficiency (POI), a condition involving the loss of ovarian endocrine function that causes long-term health effects such as infertility, heart disease, and cognitive decline in patients who have undergone gonadotoxic therapies [27,28]. We observed that, by embedding ovarian tissues within GHC grafts and implanting them together, GHCs enabled rapid assembly of functional vasculature, resulting in blood perfusion to embedded ovarian tissues within 10 days. Unexpectedly, seeded ECs did not require *in vitro* preculturing to assemble into functional vasculature inside implants; instead, preculturing the grafts was detrimental to the health of the embedded ovarian tissues. Together, this work establishes GHCs as a promising platform for endocrine tissue grafting.

## RESULTS & DISCUSSION

### 1. Interstitial fibrin-collagen supports vascular cord formation *in vitro* and perfusable network assembly *in vivo* in GHCs

We first investigated the potential for conventional MAP scaffolds to support vasculogenic assembly. Briefly, dextran, a biocompatible and minimally immunogenic polysaccharide [9,29], was functionalized with vinyl sulfones to produce a dextran vinyl sulfone (DexVS) backbone that can be crosslinked into microgels or functionalized via Michael type addition. To form microgels, monomeric DexVS solution was mixed with polyethylene glycol dithiol (P2T) crosslinker and perfused through a highly parallelized microdroplet generator [30] to rapidly generate sufficient numbers of microgels for scaffold parameterization studies (Figure 1A). P2T was selected to render microgels non-degradable, ensuring long-term graft stability in vivo. Initial microgel diameters were 66 µm (Figure 1B), matching values of previously reported MAP scaffolds composed of other synthetic polymeric materials[31,32]. To assemble microgels into larger scaffolds, microgels were interlinked with a nondegradable dithiolated peptide linker (ndVPMS).

**Figure 1:**
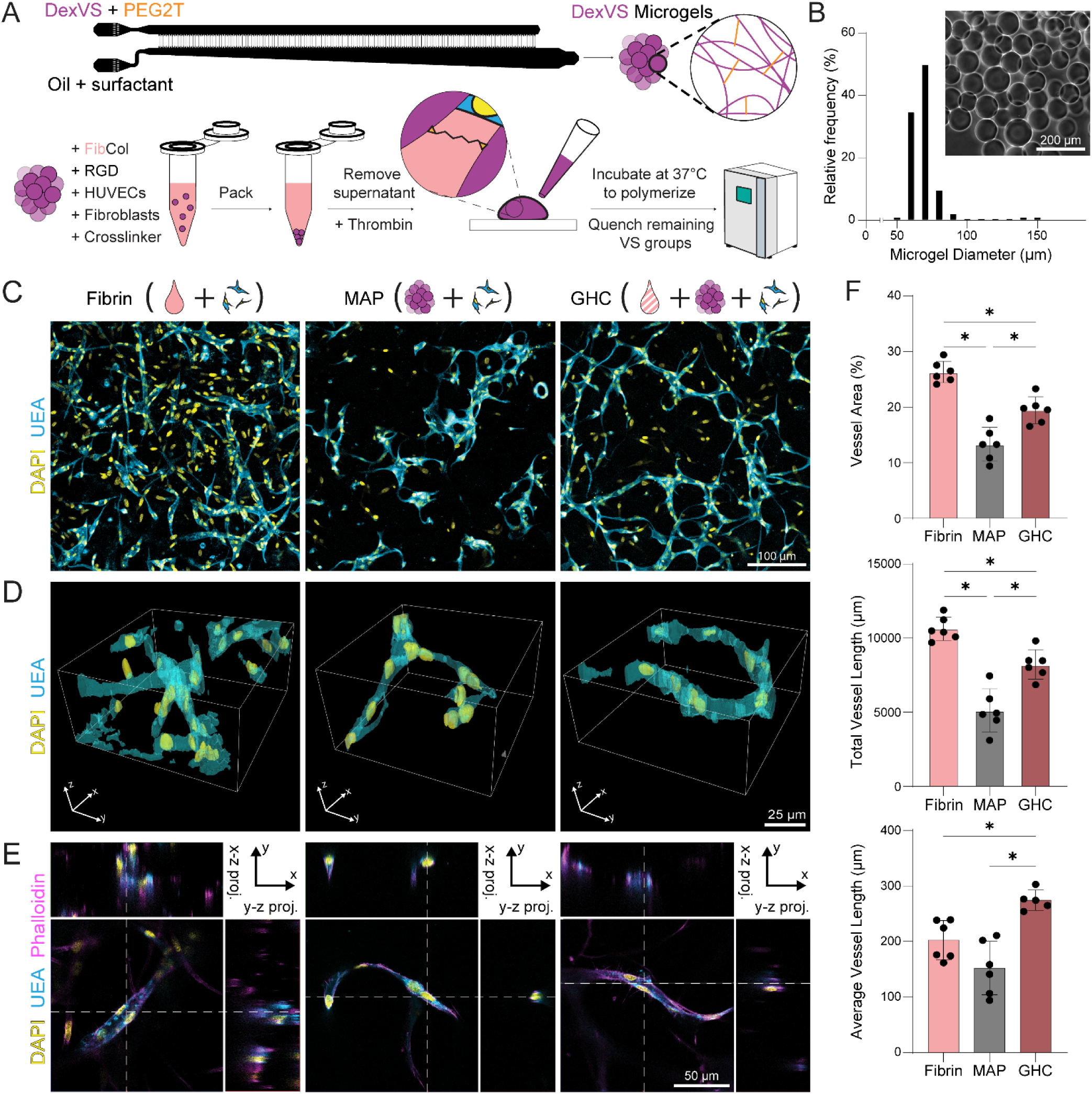
Fabrication and comparison of vascular network formation in MAP and GHC gels. (A) Fabrication schematic for GHC generation. (B) Histogram of microgel diameters. (C) Max. intensity projections of vascular networks, (D) high mag. AIVIA renderings of vascular cords, and (E) high mag. single z-slices and orthogonal views of lumenized vessels in fibrin, MAP, and GHC scaffolds. (F) Vessel area fraction, total vessel length, and average vessel length quantifications, respectively. *P < 0.05.

With the goal of driving vasculogenic assembly within MAP scaffolds, prior studies have seeded ECs within MAP scaffolds with or without support cells; however, ECs appeared to adhere to and wrap around microgels, forming flattened, surface-restricted assemblies reminiscent of EC monolayers on 2D substrates instead of cord-like networks of perfusable tubular structures, [22,26,33]. We hypothesize that these results are due to MAP scaffolds not providing adequate matrix support for ECs to assemble into perfusable vessel morphologies; EC lumen and tube formation occurs exclusively in 3D matrix settings, with lumenogenic signaling not engaging when ECs are seeded on 2D hydrogels [34–37]. To test this, interstitial pores within the MAP scaffold were filled with vasculogenic fibrin-collagen precursor solution (FibCol, 5 mg/mL fibrin and 1 mg/mL collagen)[38] to yield granular hydrogel composites (GHCs). We observed P2T causing fibrin clumping and worse vasculogenic assembly, thus motivating ndVPMS as the microgel interlinking agent for both GHCs and MAP gels (Figure S1). To compare the vasculogenic potential between MAP scaffolds and GHCs, HUVECs and NHLFs were seeded into these two scaffold formats and cultured for three days, with bulk fibrin serving as a positive control. Consistent with the literature[22,23], HUVECs in MAP scaffolds possessed morphologies atypical of vascular networks; while some ECs assembled into thin, cord-like structures, the vast majority formed monolayers on the surfaces of microgels (Figure 1C). This may arise from MAP scaffolds possessing only microgel surfaces to support cell adhesion, resulting in cell spreading on convex surfaces. As compared bulk fibrin controls, ECs in MAP scaffolds possessed smaller network area coverage, lower total network length, and shorter average vessel length (Figures 1C, F). Additionally, the cord-like structures that managed to form in MAP scaffolds appeared to be thin and non-lumenized. In contrast, cord-like structures in the fibrin gels were thick and lumenized (Figures 1D, E), supporting the hypothesis that cell interactions with a surrounding 3D matrix better supports vasculogenic assembly.

In contrast to MAP scaffolds, GHCs supported the assembly of cord-like and lumenized EC structures with comparable morphology to those observed in fibrin bulk controls (Figures 1D, E). We found FibCol infilling fostered denser vascular networks than fibrin or collagen alone; this matches results from literature [38] and was thus used for all subsequent studies (Figure S2). Network area coverage and total network length were both higher in GHC scaffolds than in MAP scaffolds, indicating more successful vasculogenic assembly, but lower than in fibrin gels. However, GHC scaffolds supported longer average vessel lengths than both MAP scaffolds and fibrin gels, indicating assembled vessels were better interconnected (Figures 1C, F). This may be an intrinsic effect of granular hydrogels: while quantifications were performed over identical overall scaffold volumes, the volume fraction that cells can inhabit varies between fibrin and GHC/MAPs. ECs are spatially confined to the interstitial space between microgels, potentially resulting in initially closer proximity of embedded cells that promotes multicellular assembly. In bulk fibrin gels, by contrast, ECs are distributed throughout the entire gel volume, resulting in initially larger intercellular distances requiring more cell migration and spreading for multicellular structures to form. Nonetheless, these results support our hypothesis that ECs in MAP scaffolds lack sufficient 3D matrix support to undergo vasculogenic assembly and demonstrate that pore-infilling with FibCol allows ECs to assemble into a cord-like network. Additionally, fibrin and collagen are also both well-known vasculogenic mediums and may provide additional vascular support beyond dimensionality, such as pro-vasculogenic fibrous cues and GF sequestration [9,16,39]. Taken together, GHCs better support vasculogenic assembly than MAP scaffolds.

Having achieved cord-like networks in granular hydrogels *in vitro*, we next implanted GHCs, MAP scaffolds, and fibrin gels to test how well each graft facilitates graft-host anastomosis and blood perfusion. Grafts were implanted into murine parametrial fat pads, an implant site that readily allows for robust graft vascularization and is used for long-term tissue engraftment, such as for islet or ovarian tissue implants [40,41]. Seven days after implantation, fibrin grafts had completely resorbed and were irretrievable, whereas MAP and GHC grafts remained integrated with the surrounding host tissue (Figures 2A, B). Ulex europaeus agglutinin 1 (UEA) staining for HUVECs revealed that GHC grafts possessed denser surviving vascular networks than MAP grafts, with higher vessel area coverage and greater total network length (Figures 2C, D). TER-119 immunostaining for murine red blood cells (mRBCs) indicated greater blood perfusion both inside human EC vascular networks and within GHC grafts overall compared to MAP grafts (Figures 2E, F). Surprisingly, explanted MAP grafts also contained cord-like structures, potentially due to the 7 days in vivo providing additional time for network assembly to occur; Zhang et al. have reported cord-like EC assembly in MAP gels after 7 days of *in vitro* culture [25]. Additionally, soluble matrix components diffusing to and localizing within the scaffold and invading stromal cell matrix deposition may help build up enough of an interstitial matrix for ECs to undergo tube formation within MAP gels. However, GHCs possessed more vascular cords than MAP gels after 3 days of preculture, resulting in a greater number of vessels supporting host blood flow. Taken together, pre-vascularized GHCs formed denser and more functional vascular networks than pre-vascularized MAP gels, resulting in better vascular integration with host vasculature.

**Figure 2:**
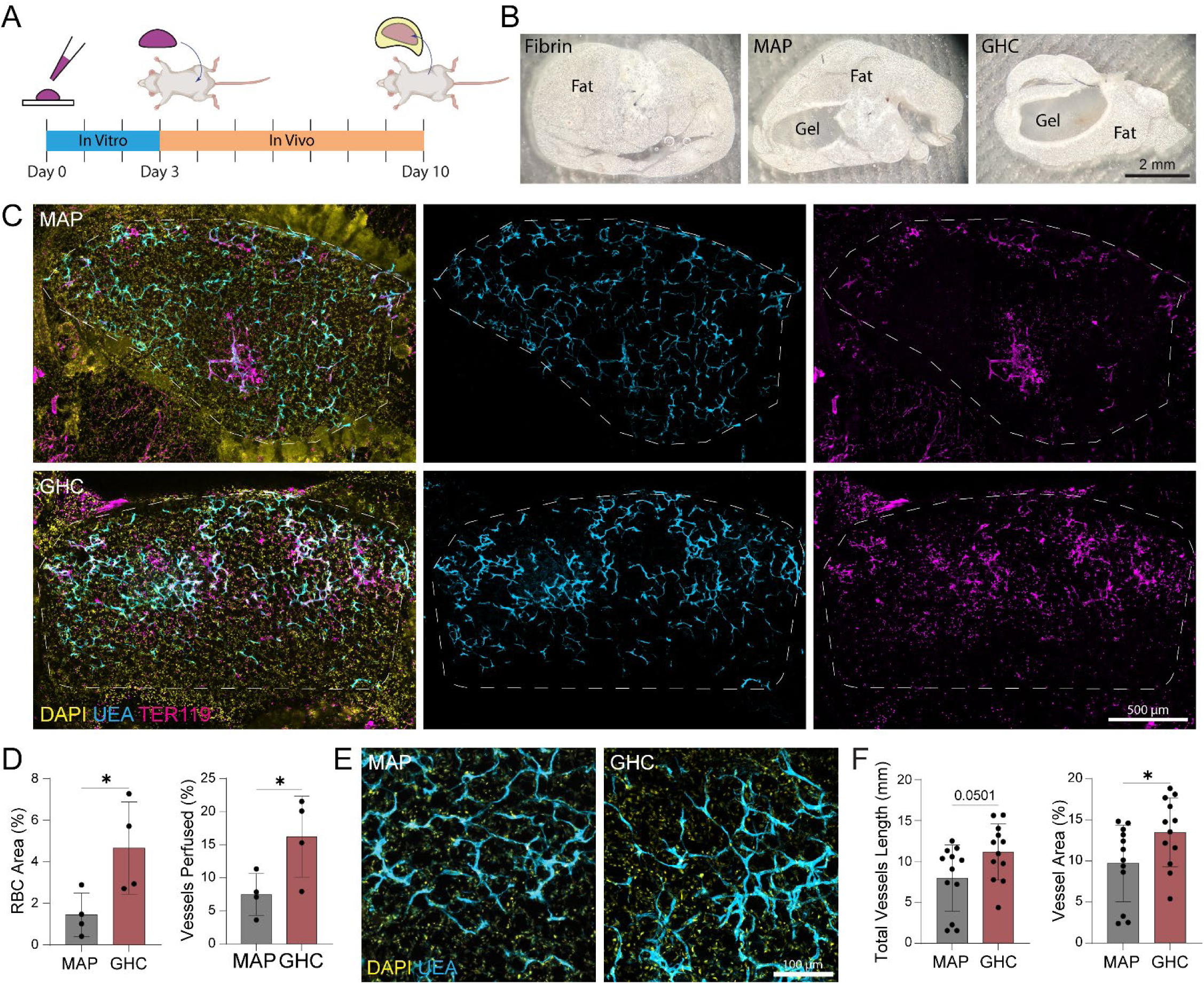
Implantation and In Vivo comparison of vascular integration in MAP and GHC gels. (A) Timeline of graft culture and implantation. (B) Cross-sectional slices of processed explants. (C) Confocal max. intensity projections of MAP and GHC graft slices. (D) Graft RBC amount and perfused vessel fractions quantifications, respectively. (E) Fluorescent images of HUVECs in explanted grafts. (F) Vessel network area and total vessel length quantifications, respectively. *P < 0.05.

### 2. RGD functionalization and microgel interlinking enhance vasculogenic assembly in GHCs

In typical MAP scaffolds, microgels are functionalized with RGD and interlinked to generate a handleable graft that supports cell adhesion. However, with the addition of FibCol to the pore space, embedded or invading host cells can attach to integrin binding motifs including RGD and GFOGER presented by FibCol. As such, we wondered whether microgel functionalization with RGD was required for vasculogenic assembly. To examine this, DexVS microgels were functionalized with varying concentrations of RGD (0 - 4.25 mM) prior to GHC fabrication. After three days of culture, HUVECs in all conditions formed cord-like networks. Vessel network coverage and average vessel length increased with increasing RGD concentration, while total vessel length had the largest difference between 0 and 1 mM (Figure S3). These results demonstrate that microgel functionalization with RGD improves vascular network assembly, suggesting that integrin-mediated interactions are beneficial to such process. Given that integrins facilitate adherent cell spreading and cell migration, RGD functionalization may be allowing ECs to migrate quicker, resulting in the increases in vessel area coverage and average vessel length. Since the 4.25mM RGD condition produced the vascular networks with largest vessel area coverage and longest average vessel length, this concentration was used for all subsequent experiments.

The inclusion of FibCol between microgels not only enables integrin-mediated cell attachment and supports lumen formation, but also acts to physically bind microgels together, enabling the creation of a handleable graft without the requirement for microgel interlinking (Figure S4). However, engraftment environments are highly proteolytic and may degrade the FibCol away prior to successful vasculogenic assembly. To test whether interlinking microgels with covalent crosslinks was necessary for perfusable vascular network formation, GHC grafts were assembled with or without the ndVPMS peptide linker and implanted as in studies described earlier. Assessment of explants indicated that interlinked GHC grafts contained higher vascular network coverage, greater total vessel length, and longer average vessel length along with a greater number of mRBC-perfused vessels as compared to GHC grafts where microgels were not covalently interlinked (Figure S5). This difference may be due to the permanence of the microgel binding; FibCol degradation weakens its ability to physically bind microgel together over time, while ndVPMS interlinking remains stable throughout implantation. Given that ndVPMS interlinking resulted in more mRBC-perfused vessels, this method was used to construct GHC grafts for all subsequent studies.

### 3. Microgel diameter influences assembled microvessel diameters, density, and resulting graft perfusability

Previous studies focused on MAP scaffolds have explored a range of microgel diameters, finding dispersed cells in MAPs composed of larger microgels (ø = 170-200 μm) percolate via gravity through scaffold pores, resulting in a nonuniform distribution of embedded cells [22,42]. The rapid gelation of FibCol in our GHCs prevents this from occurring, allowing for creation of grafts composed of larger microgels while maintaining a uniform distribution of encapsulated cells. Based on our previous finding that angiogenic sprouts greater than 11 μm in diameter reliably form lumens [43], larger interstitial spaces between microgels may allow for the formation of thicker vascular cords with greater potential for lumenization required for forming perfusable vascular networks. To test whether microgel diameter and resulting interstitial geometry influences the diameters of formed vascular cords, we began by characterizing interstitial geometry as a function of microgel diameter. GHCs were formed with fluorescently tagged microgels ranging in diameters 83-170 μm (Figure 3A, B); channel heights in the microdroplet generator chip were increased to create microgels with larger diameters [30]. Interstitial area fraction was analyzed by binarizing 2D slices from z-stack images. Despite differences in microgel size, average values remained consistent at ∼25% of scaffold cross-sectional area (Figure 3D). This 2D measurement has been previously shown to return equivalent results to volumetric analyses [44]; as such, GHCs across all microgel sizes maintain a volumetric interstitial fraction of ∼25%, consistent with the theoretical ∼26% void fraction predicted computationally for ideally packed solid spheres [45]. Interstitial geometry was quantified by the diameter of the largest circle fit between microgels (Figure 3C). Average fitted circle size and the fraction of fitted circles >11 μm increased with microgel size, while number of fitted circles per slice decreased with microgel size (Figure 3F). Together, these results demonstrate that microgel diameter is an effective handle for tuning interstitial architecture, modulating both interstitial space size and distribution within the scaffold. Critically, the majority of fitted circles in GHCs made from medium (121 ± 5.6 μm) and large (155 ± 17.5 μm) microgels exceeded the 11 μm lumenization threshold, compared to only a quarter in scaffolds formed from small (66 ± 5.2 μm) microgels, motivating further investigation into how these geometric differences influence vasculogenic assembly.

**Figure 3:**
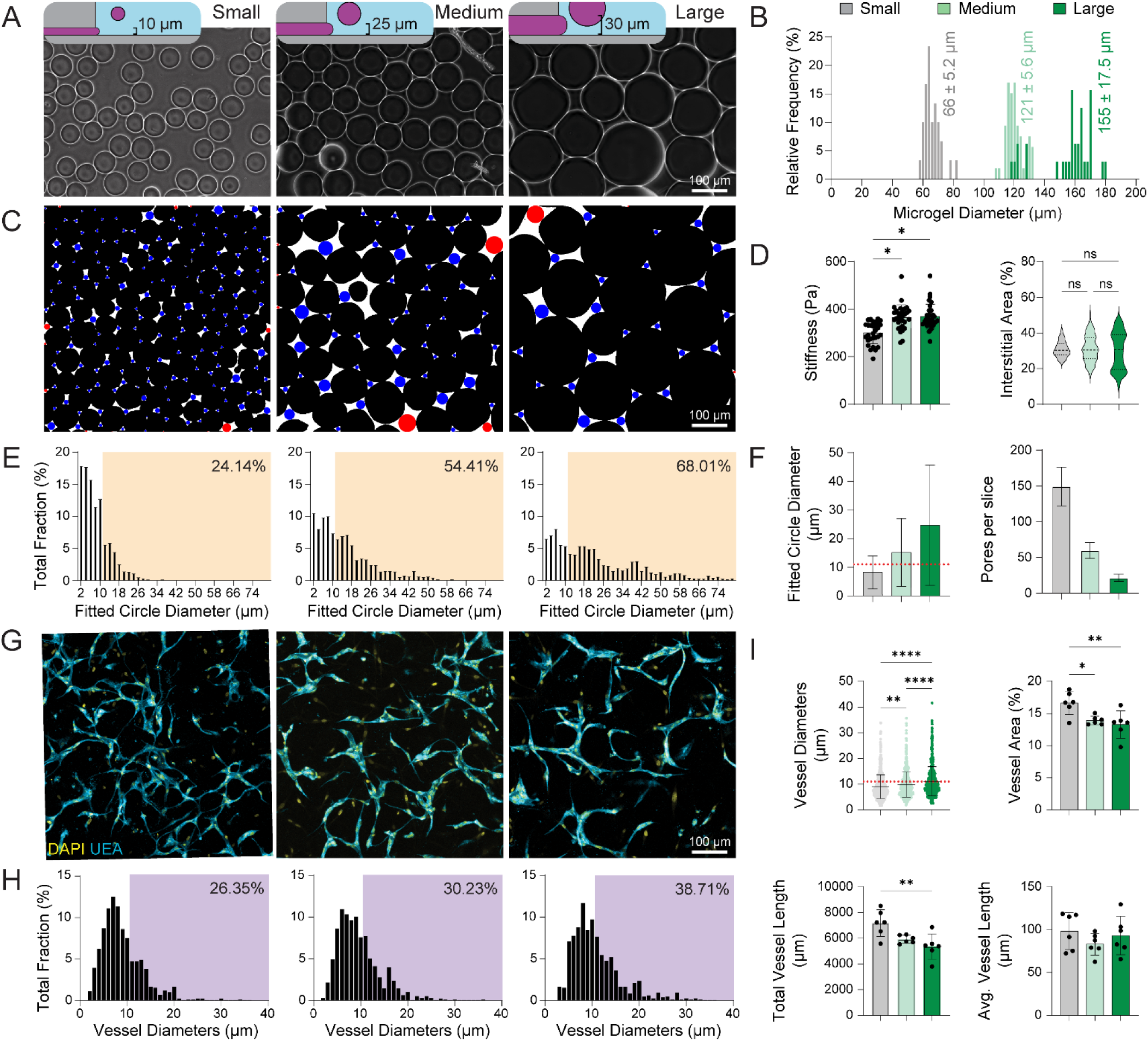
Microgel size influences assembled vascular network properties. (A) Brightfield images of microgel populations with different diameters. (B) Histogram of microgel diameters from all three populations. (C) Image analysis to quantify pore size by the largest circle to fit in each pore. Red circles were omitted due to border artifacts. (D) interstitial area quantification. (E) Histogram of fitted circle diameters and percentage larger than or equal to 11 μm. (F) Pore size and quantity quantifications. (G) Confocal fluorescent images of vascular networks assembled within GHCs of different microgel sizes. (H) Histogram of vessel diameters and percentage of measurements larger than or equal to 11 μm. (I) Vessel diameter, vessel network area, total vessel length, and average vessel length quantifications, respectively. *P < 0.05.

To test whether the geometry of interstitial space alters vascular cord lumenization, we morphometrically analyzed vascular networks after 3 days of assembly in GHCs as a function of microgel diameter (Figure 3G). Increasing microgel size corresponded to both larger cord diameters and a higher percentage of cords possessing >11 μm diameters (Figure 3H, I). Interestingly, network area coverage and total vessel length decreased with increasing interstitial space size, while average vessel length remained unaffected (Figure 3I). This suggests that interstitial geometry has minimal influence on vascular network interconnectivity but plays a substantial role in determining overall network density, an outcome likely reflecting differences in interstitial architecture arising from varied microgel sizes. Since overall interstitial volume is conserved regardless of microgel size, increasing interstitial space size necessarily reduces interstitial space frequency to compensate (Figure 3D, F). Thus, though larger microgels allow for thicker vascular cords within larger interstitial spaces, they also reduce vascular network density by decreasing the number of interstitial spaces available for vessel assembly. Given that thicker cords are more likely to be lumenized [43], these data demonstrate that GHCs formed from larger microgels guide ECs to assemble more sparsely distributed networks but with greater potential for lumenization.

Given this potential tradeoff between vascular network density and individual cord diameter, we next tested *in vivo* to observe how these differences in vascular architecture affected anastomosis with host vasculature. GHC grafts spanning the same range of microgel sizes were pre-cultured for 3 days and then implanted into the parametrial fat pads of female mice. Despite previous *in vitro* findings favoring smaller microgels for network density, medium GHC grafts possessed the densest vascular networks of the three conditions *in vivo*, while small and large GHC grafts formed vascular networks of similar densities. (Figure 4A, B). This suggests that both vessel thickness and network density contribute to graft vascularization, with medium GHCs achieving a balance between the two. Evaluation of mRBC presence within these grafts revealed that, though all constructs had similar percentages of their vascular networks perfused, the medium GHC grafts contained more mRBCs than the small GHC grafts and were more consistent than the large GHC grafts (Fig 4C, D). This indicates that larger microgels, via larger pores, promote more perfusable vasculature. However, this effect plateaus at the largest microgel size, with large GHC grafts showing high variability and performing worse than medium GHC grafts on average. Given that networks in large GHC grafts were sparser than the other two conditions, host vessels may have had a harder time finding and connecting to EC networks in large GHC grafts reliably, resulting in the binary outcomes in large GHC grafts. Taken together, these data demonstrate that GHC pore size regulates both vascular network density and vessel thickness, and that balancing these two parameters maximizes graft anastomosis and perfusability.

**Figure 4:**
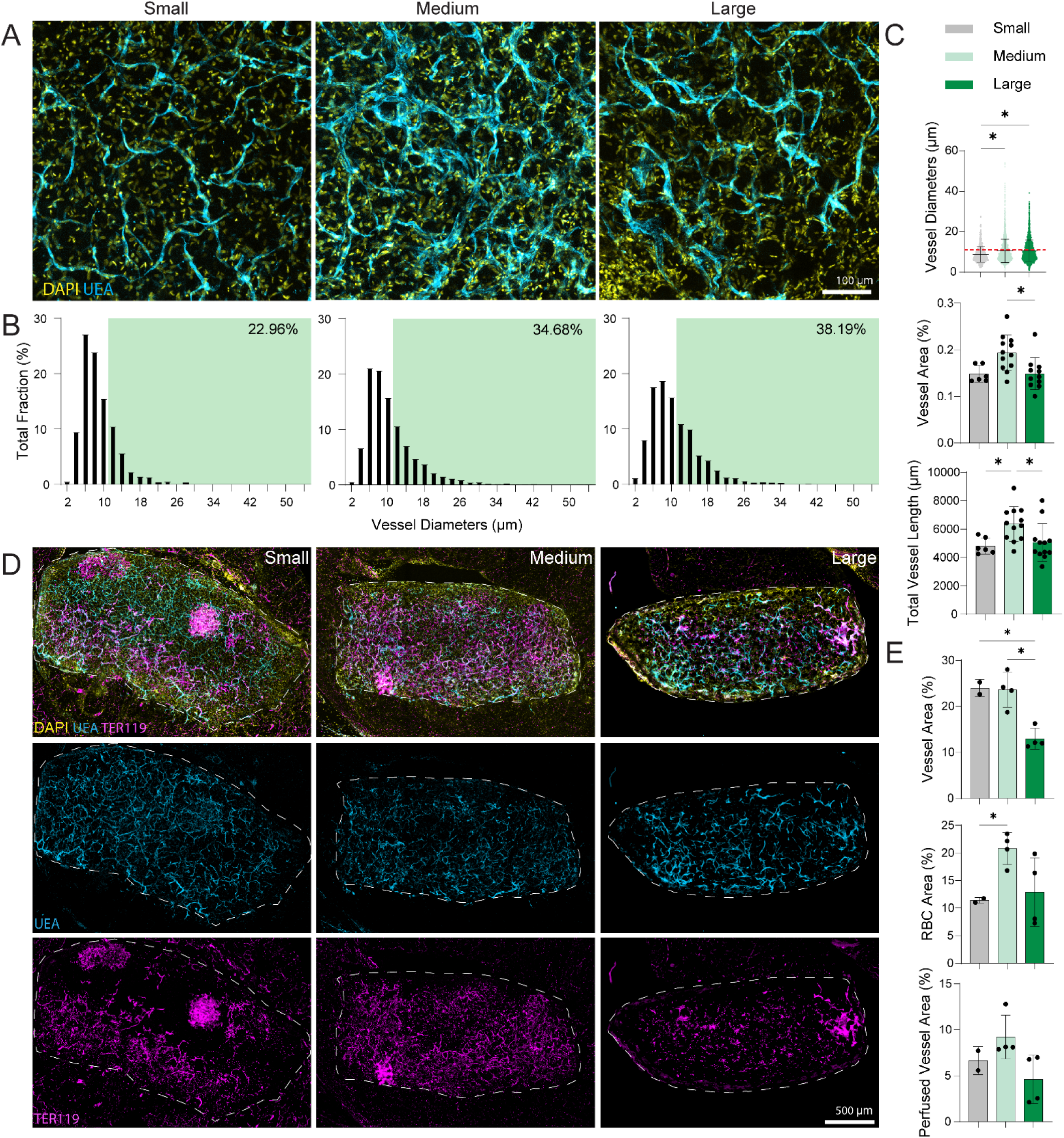
Medium microgels best support vascular integration of GHCs. (A) Confocal fluorescent images of HUVEC networks in GHC grafts as a function of microgel size. (B) Histogram of vessel diameters and percentage of measurements larger than or equal to 11 μm. (C) Vessel diameter, vessel network area, and total vessel length quantifications, respectively. (D) Confocal max. intensity projections of GHC graft slices as a function of microgel size. (E) Graft vessel area, RBC area, and perfused vessel area quantifications, respectively. *P < 0.05.

### 4. GHCs enable rapid vascular assembly, integration, and perfusion of implanted ovarian tissues

Having optimized GHCs to support perfusable vasculature *in vivo*, we next tested whether this platform could promote the vascular integration of embedded parenchymal tissue with therapeutic relevance. Ovarian tissues are a challenging tissue to engraft given their high metabolic demands and sensitivity to external perturbations such as non-ovarian growth factors, heat stress, and mechanical fragmentation [46,47]. Clinically, ovarian tissue grafting is performed as part of ovarian tissue cryopreservation and transplantation (OTCT), a clinical intervention seeking to preserve endocrine function and fertility in patients suffering from primary ovarian insufficiency (POI), the premature loss of ovarian function arising from gonadotoxic cancer treatments [27]. POI has been shown to cause long-term health complications, including cardiovascular disease, osteoporosis, mental health issues, and cognitive decline [28]. OTCT involves extracting and preserving healthy ovarian tissue prior to the gonadotoxic treatments, followed by implantation after cancer remission [48]. However, OTCT has had limited success, with the duration of tissue function varying between 0.7 and 5 years, likely due to slow tissue revascularization and resulting ischemic injury [49]. Indeed, animal studies indicate 50-90% follicular loss upon ovarian tissue implantation which severely compromises the longevity of restored endocrine function [50–52]. As such, ovarian tissues are a demanding and clinically relevant model for evaluating whether GHCs can accelerate parenchymal tissue revascularization and survival. However, *in vitro* culture of ovarian tissue results in rapid follicle loss, precluding the three-day culture period provided for vascular assembly in the studies above. As such, we asked whether a pre-implantation culture period was necessary for vasculogenic assembly in ovarian tissue-laden GHC grafts.

To test this, we embedded human ovarian tissue (∼1 mm^3^) into GHCs and implanted grafts into murine parametrial fat pads directly after graft formation (non-cultured) or after 3 days of culture in EC media (cultured). To isolate the effects of supporting vasculature from ovarian tissue culture, we tested grafts with or without embedded HUVECs and HLFs (Figure 5A). TER119 staining of explanted grafts revealed mRBC perfusion across both the GHC-fat and GHC-ovarian tissue interfaces, demonstrating successful vascular formation, anastomosis, and reperfusion (Figure 5B).

**Figure 5:**
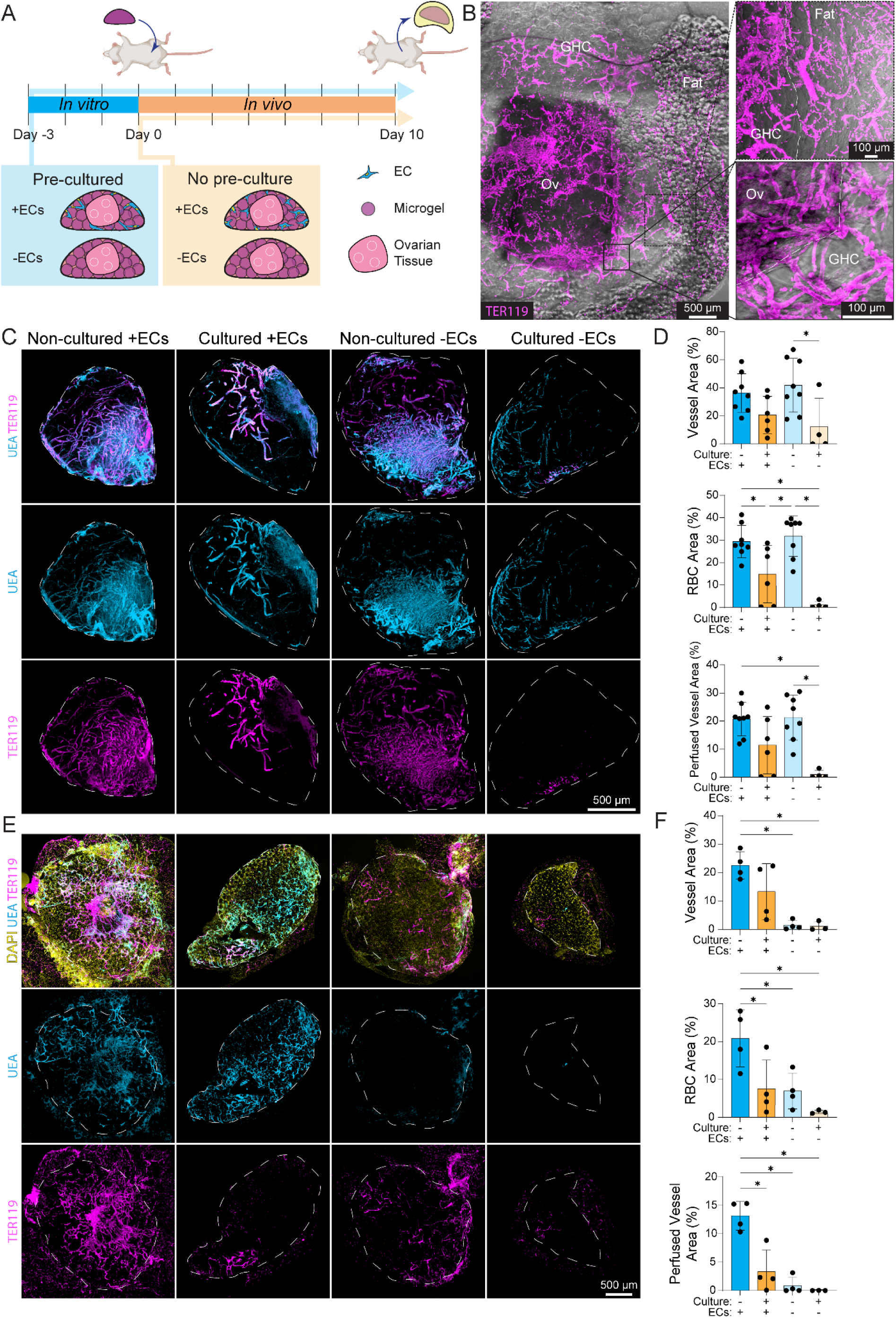
GHCs facilitate blood circulation between embedded ovarian tissues and host vasculature. (A) Timeline and overview of ovarian GHC graft culture and implantation. (B) Brightfield & fluorescent image of processed explant; mouse RBC is perfused through the fat-GHC and GHC-ovarian tissue interfaces. (C) Cross-sectional slices of ovarian tissues within implants. (D) Ovarian vessel, RBC, and perfused vessel quantifications, respectively. (E) Cross-sectional slices of the GHC portions of implants. (F) Inter-GHC vessel, RBC, and perfused vessel quantifications, respectively.

We next examined the vascular networks within the embedded ovarian tissues to gauge tissue health. We found that acellular cultured grafts performed the worst, with surviving ovarian vasculatures appearing thin, sparse, and blebby. This was reflected quantitatively, where ovarian tissue in acellular cultured grafts had the smallest vascular area fraction and minimal mRBCs as identified by TER-119 immunostaining within the tissue. In contrast, seeded cultured grafts resulted in markedly improved survival of ovarian vasculature, with thicker vessels perfused with mRBCs. These results suggest that supporting engineered vasculature is beneficial to the survival of ovarian vasculature, potentially by accelerating anastomosis, allowing for faster reperfusion and reduced ischemia. Alternatively, biochemical signals from engineered vasculature may provide survival signals. However, despite evidence of perfusion, ovarian vascular networks in seeded cultured grafts remained sparse in comparison to non-cultured grafts. Both seeded and acellular non-cultured grafts possessed dense, perfused ovarian vasculature, with larger vascular area fractions, mRBC area fractions, and greater UEA-TER119 co-localization than cultured grafts (Figure 5C, D). Overall, these outcomes indicate that pre-culture is detrimental to the health of the vascular networks within ovarian tissue. While it is possible that suitable culture conditions could be identified for ovarian tissue, direct implantation proved effective and is translationally more feasible. Unexpectedly, the absence of engineered vasculature did not affect ovarian vascular survival, as acellular non-cultured grafts performed comparably in all metrics (Fig. 5D). To better understand this unexpected finding, we next examined the surrounding engineered vasculature within the GHC.

Examining engineered vascular networks within GHC regions of the grafts, we found not surprisingly that seeded grafts contained significantly more vasculature than acellular grafts. Non-cultured samples contained more mRBCs than their cultured counterparts but, curiously, seeded cultured grafts possessed comparable amounts of mRBCs to acellular non-cultured grafts (Figure 5E, F). mRBCs in the acellular non-cultured grafts are likely invading and perfused mouse vessels that anastomosed to the ovarian tissues, accounting for the mRBCs observed within the acellular non-cultured ovarian vasculature. However, for samples that possessed both healthy ovarian tissue and engineered vasculature, we saw extensive, perfused vasculature bridging the ovarian and surrounding host tissues. Seeded non-cultured grafts had the largest vessel area, most RBCs, and greatest UEA-TER119 colocalized area, indicating the most extensive and most perfused vascular network of all conditions. Compared to the seeded cultured condition, seeded non-cultured grafts had comparable vascular networks but significantly more mRBCs and perfused vasculature. This suggests that the pre-culture period negatively affected engineered vascular functionality, possibly due to growth factor signaling from unhealthy ovarian tissue. Based on these findings, graft pre-culture is unnecessary for vascular formation, as GHC-seeded HUVECs and HLFs can form functional vasculature without pre-assembly. Compared to acellular non-cultured grafts, seeded non-cultured grafts had substantially more total and perfused vasculature, demonstrating that engineered vasculature within GHCs enhances overall graft vascularization and inosculation. This contrasts with the ovarian vasculature analysis above, where engineered vasculature did not affect ovarian vascular survival or function. However, given that a more extensive vascular network can support greater volumetric blood flow, ovarian tissues in seeded non-cultured grafts likely receive greater perfusion than those in acellular non-cultured grafts. Overall, these experiments demonstrate the ability of GHCs to foster robust vascular assembly and network integration with both embedded and host tissues. Through these capabilities, GHCs connected implanted ovarian tissues to host vasculature within 10 days, a marked improvement over the 3 weeks previously reported for MAP gels.

## CONCLUSION

In this work, we enabled cord-like EC assembly in MAP gels by introducing fibrin and collagen into the pore space. To demonstrate the advantage of assembling cord-like vascular networks, we implanted both MAP and GHC grafts and found GHCs to more readily host perfusable vasculature. To enhance vasculogenic assembly in GHCs, we investigated microgel RGD functionalization and interlinking and found both modifications to encourage network formation. Additionally, we investigated microgel size and its effects on perfusable vascular formation and found scaffolds made from microgels with diameters around 120 µm supported the most perfusable vasculature. Through incorporation of ovarian tissues into GHC grafts, we demonstrated that GHCs improve vascular integration of embedded tissues and that vascular assembly prior to implantation is unnecessary. Compared to established ovarian grafting methods, this vascularization approach yields more robust tissue reperfusion over shorter time periods. We anticipate this biomaterial design will have broad applications in vascularizing tissue grafts of other parenchymal tissues.

## EXPERIMENTAL

### Cell Culture and biological reagents

Human umbilical vein endothelial cells (HUVECs) were purchased from Lonza and cultured in endothelial growth medium (EGM-2; Lonza, Basel, Switzerland) supplemented with 1% penicillin-streptomycin-fungizone (Gibco, Waltham, MA). Human lung fibroblasts (HLFs) (University of Michigan Central Biorepository) were cultured in DMEM (Gibco, Grand Island, NY) containing 1% penicillin/streptomycin, L-glutamine and 10% fetal bovine serum (Atlanta Biologics, Flowery Branch, GA). All cells were cultured at 37 °C and 5 % CO_2_. HUVECs and HLFs were used prior to passage 8 and 10, respectively.

### Synthesis of dextran vinyl sulfone

Dextran was functionalized with divinyl sulfone following a previously described procedure [53]. Briefly, dextran (5 g) was dissolved into 250 mL of sodium hydroxide (100 mM) solution on a stir plate at 1500 rpm before the addition of divinyl sulfone (12.5 mL; TCI America, Portland, OR). The reaction proceeded for 3.5 min. before termination by the addition of 2.5 mL hydrochloric acid (12 M). The product was dialyzed against Milli-Q water for 3 days and then lyophilized to produce a dry product. Reaction products were analyzed with 1H NMR.

### Microgel Fabrication and characterization

DexVS microgels were generated on a high-throughput microfluidic droplet generating device adapted from De Rutte et al. [30]. The device was designed in AutoCAD and a master mold was fabricated using SU-8 negative photoresist (Kayaku, Westborough, MA) and standard photolithography. Polydimethylsiloxane (PDMS) (Dow, Midland, MI) was prepared at a 1:10 cross-linker:base ratio and cast onto the silicon/SU-8 mold to generate microfluidic devices, which were then cleaned, bonded to glass via plasma etching, and silane-treated. Shelf heights of 10, 25, and 30 μm were used to generate the small, medium, and larger microgel populations. Two aqueous solutions were prepared by separately dissolving DexVS (19% functionalization) and the crosslinker poly(ethylene glycol) dithiol (P2T) in PBS with 50 mM HEPES buffer.

A non-aqueous phase was prepared by adding 1.5 wt./v% perfluoropolyethylene (Ran Biotechnologies, Beverly, MA) to HFE-7500 (3M, St. Paul, MN), a fluorinated engineered fluid. A syringe pump was used to flow the aqueous and non-aqueous solutions through the microfluidic droplet generating device at 1.0 and 2.0 mL/hr., respectively. Aqueous solutions were first perfused through a custom PDMS microfluidic mixing device to mix the P2T crosslinker with DexVS. The resulting mixture then traveled through Tygon tubing to the droplet generating device to generate monodisperse aqueous-in-non-aqueous droplets.

The emulsion was collected and then broken by the addition of PBS and 30 v/v% perfluorooctanol (PFO) (Alfa Aesar, Haverhill, MA). The non-aqueous phase was removed from collected microgels via centrifugation for 2 min. at 3000 g, washed 3x with PBS and hexanes simultaneously, then washed 3x with 0.1wt/v% bovine serum albumin (BSA) solution. Washed microgels were stored in 0.1wt/v% BSA solution with 4x Primocin.

### Granular hydrogel formation, culture, and characterization

Microgels were functionalized with 4.25 mM RGD, then suspended at a 1:2 volumetric ratio with 2 million/mL HUVECs, 500k/mL HLFs, and 5mM ndVPMS dithiol peptide linker [54] in 50mM HEPES solution or 5 mg/1 mg/mL fibrinogen-collagen solution to form microporous annealed particle (MAP) or granular hydrogel composite (GHC) gels, respectively. Solutions were homogenized via pipetting and centrifuged at 500x g for 2 minutes to pack microgels and cells, concentrating HUVECs and HLFs between microgels to final concentrations of 4 million/mL and 1 million/mL, respectively. Supernatant was then removed, and 1 U/mL thrombin was mixed into the packed GHC samples. Microgel/cells were then pipetted into custom fabricated 4 mm diameter PDMS gaskets that held 20 μL of volume, resulting in cylindrical-shaped constructs which were allowed to anneal and polymerize for 30 min. at 37 °C. Afterwards, constructs were submerged for 30 min. in 50mM HEPES solution with 1x M199 and 24mM cysteine to quench unreacted vinyl sulfone groups. Constructs were subsequently washed 2x with PBS and then cultured in vasculogenic media consisting of EGM2 (Lonza) supplemented with 5 vol% FBS, 50 ng/mL VEGF, and 25 ng/mL PMA. Media was replenished every 24 hrs. For linker concentration screening, GHCs were formed with 0, 5, 10, and 15mM of D-VPMS. Constructs were imaged, then incubated in 10 mg/ml nattokinase (JBSL-USA, Walnut Creek, CA) and 1 mg/ml Dispase II (Sigma-Aldrich) for 12 hrs. to completely degrade away fibrin and collagen, respectively.

Fibrin hydrogels (5 mg/mL fibrinogen, 1 U/mL thrombin) were seeded with 4 million/mL HUVECs and 1 million/mL HLFs. Briefly, a fibrinogen precursor solution was prepared by dissolving fibrinogen from bovine plasma in PBS at twice the final concentration. A thrombin precursor solution was simultaneously prepared at 2 U mL-1 resuspended in EGM-2 consisting of cells at twice the final concentrations; for the interlinking agent compatibility study, 12.25 mM of P2T and ndVPMS were also added. These two precursor solutions were then mixed at a 1:1 ratio, and 20 μL of this solution was transferred into a PDMS mold with 4 mm diameter gasket for incubation at 37 ◦C for 30 min. Fibrin gels were then cultured with vasculogenic media as outlined above.

### Fluorescent staining and microscopy

Hydrogels were fixed with 4% paraformaldehyde for 1 hr. at room temperature and permeabilized in PBS containing Triton X-100 (0.5 v/v%), sucrose (10 w/v%), and magnesium chloride (0.6%w/v). To visualize actin cytoskeleton and nuclei, samples were stained with phalloidin and DAPI (Sigma-Aldrich) for 1 hr. at room temperature. To visualize human endothelial cells, samples were stained with Ulex Europaeus Agglutinin I (UEA-I) (1:200, Vector Labs, Burlingame, CA) for 1 hr. at room temperature. Fluorescent images were captured on a Zeiss LSM800 laser scanning confocal microscope. For vascular network analysis, z-stacks were acquired with a 10x objective. High-resolution images were similarly acquired with a 40x objective. All images are presented as maximum intensity projections unless otherwise specified.

### Image analysis

Image stacks were transformed into maximum intensity projections for analysis. Morphological properties of vascular networks were quantified using Angiotool, while vessel diameters were quantified using REAVER [55,56]. For vessel area, total network length and average segment length quantification, vessel-defining parameters specified in Zudaire et al., 2011 were used; a vessel is defined as a segment between two branching points or a branching point and an endpoint [55]. Vessel area percentage is the area of the segmented vessels divided by the area of the images. The total length is the sum of the Euclidean distances between the pixels of all the vessels in the image, while the average vessel length (average segment length) is the mean length of all the vessels in the image (i.e. total length divided by the total number of vessels in the image). For vessel diameter quantifications, parameters specified in Corliss et al. [56] were used.

To characterize interstitial spaces inside GHC, microgels were tagged with custom thiol-ended AlexaFluor647 peptides, formed into GHC gels, and imaged as previously described. Z-stacks were input into a custom MATLAB code where each slice was binarized to define microgel vs. interstitial space. Interstitial fraction was defined as the total area of all interstitial spaces divided by the total area of input slice. For each interstitial space, the code determined the largest radius of a circle that could fit inside said space; circles fitted onto the edge of images were excluded from this calculation.

### Ovarian tissue procurement and encapsulation

As previously described [41,57,58], whole human ovaries were obtained from deceased donors for non-clinical research through the International Institute for the Advancement of Medicine (IIAM) and associated Organ Procurement Organizations (OPOs). IIAM and the associated OPOs comply with state Uniform Anatomical Gift Acts (UAGA) and are certified and regulated by the Centers for Medicare and Medicaid Services (CMS). These OPOs are members of the Organ Procurement and Transplantation Network (OPTN) and the United Network for Organ Sharing (UNOS) and operate under standards established by the Association of Organ Procurement Organizations (AOPO) and UNOS. Informed, written consent was obtained from the deceased donors’ families prior to tissue procurement for the tissue used in this study. A biomaterial transfer agreement is in place between IIAM and the University of Michigan that restricts the use of the tissue to pre-clinical research not involving the fertilization of gametes. The use of deceased donor ovarian tissue in this research is categorized as “not regulated” per 45 CFR 46.102 and the “Common Rule” as it does not involve human subjects and therefore complies with the University of Michigan Institutional Review Board (IRB) requirements.

Ovarian tissues were collected and prepared following previously described procedures [41,58]. Briefly, outer cortex layers were cut from donor tissues into approximately 1 mm x 10 mm x 10 mm tissue squares and cryopreserved for storage. Prior to GHC formation, cortex squares are thawed and further cut into cubes approximately 1 mm in length and width in thawing solution. To encapsulate tissues in GHCs, GHC precursor slurries were pipetted into PDMS gaskets as described above and ovarian tissue cubes were inserted immediately afterwards. Tissue-embedded GHCs were then incubated at 37 °C for 20 minutes to polymerize and cysteine quenched as described above. Non-cultured GHC samples were implanted right after cysteine quenching, while cultured samples were incubated with vasculogenic media for 3 days prior to implantation.

### In vivo studies and staining

Constructs for mouse implantation studies were generated as described above and removed from PDMS gaskets immediately before implantation. Prior to implantation, cells were cultured for 3 days in vasculogenic media described in the granular hydrogel formation, culture, and characterization section. All animal procedures in this work were performed in accordance with the protocol approved by the Institutional Animal Care and Use Committee (IACUC) at the University of Michigan (PRO00011284). Female NSG mice (strain 005557, The Jackson Laboratory, Bar Harbor, ME, USA) 10–11 weeks old were anesthetized by isoflurane and treated with Carprofen (5 mg/kg, subcutaneously, Rimadyl, Zoetis) for analgesia. The intraperitoneal space and the parametrial fat pads were exposed through a midline incision and secured using an abdominal retractor. For the ovarian vascularization study, mice were also ovariectomized prior to implantation by cauterizing the uterine horns directly below the ovary and cutting off the ovaries with microscissors. Hydrogels (one on each side) were wrapped within parametrial fat tissue, sutured in place with 10–0 Nylon sutures, and the fat tissue and constituent grafts were returned to the abdominal cavity. The muscle and skin layers of the abdominal wall were closed with 5/0 absorbable sutures (AD Surgical). All conditions consisted of 4 sample repeats, with 2 samples implanted per mouse. The mice were allowed to recover in a clean, warmed cage and received another dose of Carprofen 24 h post-recovery or as needed.

Whole fat pads were retrieved after 7 days or 10 days of implantation. Mice were anesthetized as described above. A longitudinal midline incision was made in the abdominal wall, and whole fat pads containing hydrogels implants excised and fixed with 4% paraformaldehyde solution over night at 4 °C. Tissues were washed three with in PBS for 1 hour each and stored in 0.1% azide in PBS at 4 °C.

Fixed tissues were embedded in 1.6-3.2 wt% low melting agarose in PBS and sectioned via vibratome. Sections with implanted hydrogels were permeabilized overnight at 4 °C and blocked with 5% (w/v) goat serum. For mouse erythroid cell visualization, sections were stained with the primary antibody TER-119 (1:200, Thermo Fisher Scientific, Waltham, MA). Sections were washed with PBS and incubated in goat anti-rat Alexa Fluor-conjugated secondary (1:1000, Thermo Fisher Scientific, Waltham, MA) overnight. For human endothelial cell visualization, sections were stained with UEA-I (1:200, Vector Labs, Burlingame, CA) and DAPI (1:500, Sigma-Aldrich). After staining, tissue sections were mounted with Fluoromount (Thermo Fisher Scientific, Waltham, MA).

Tissues sections were imaged on a Zeiss LSM800 laser scanning confocal microscope. Vascular network analysis was performed as previously mentioned, while custom-built MATLAB scripts were used to perform blood analysis on implants. Z-stacks (50-200 µm) of entire implants in tissue sections were acquired with an 5x objective. Maximum intensity projections of the UEA and TER119 channels were input into a custom MATLAB code which separately thresholded each channel to remove background. A user-drawn ROI covering the implant was used to separate the implant from surrounding tissues. Vessel Area percentage was defined as number of pixels positive for UEA inside ROI over total number of pixels inside ROI. RBC area percentage was defined as number of pixels positive for TER119 inside the ROI over total number of pixels inside ROI. Perfused vessel percentage was defined as number of pixels positive for both TER119 and UEA over total number of pixels positive for UEA. Perfused vessel area percentage was defined as number of pixels positive for both TER119 and UEA over total number of pixels inside ROI.

### Statistical analysis

Statistical significance was determined by one-way analysis of variance (ANOVA) with post-hoc analysis (Tukey test), Kruskal-Wallis, or Student’s t-test where appropriate. For all studies, significance was indicated by p < 0.05. Sample size is indicated within corresponding figure legends, and all data are presented as means ± standard deviations.

## AUTHOR CONTRIBUTIONS

M.M.H, D.I.P, A.S., and B.M.B designed the experiments. M.M.H, C.L., G.A.G., and F.S.M. contributed to microgel fabrication and *in vitro* experiments and data analysis. M.M.H., D.S., C.L., K.L, Y.Z., and A.S. contributed to *in vivo* experiments and data analysis. R.N.K. provided initial MATLAB code for interstitial space analysis. M.M.H. and B.M.B wrote the manuscript. Y.Z. and F.S.M. edited the manuscript. All authors reviewed the manuscript.

## DECLARATION OF CDOMPETING INTEREST

The authors declare that they have no known competing financial interests or personal relationships that could have appeared to influence the work reported in this paper.

## ACKNOWLEDGEMENTS

M.M.H. acknowledges financial support from the National Institute of Dental & Craniofacial Research of the National Institutes of Health Tissue Engineering at Michigan T32 Training Grant (T32DE00705745). B.M.B. acknowledges financial support from NIH (RO1EB030474).

