## Supplemental Data for "Filling the Void: Rapid Revascularization via Vasculogenic Assembly in Semi-synthetic Granular Hydrogel Grafts"

Granular hydrogel

Graft Revascularization

Vasculogenesis

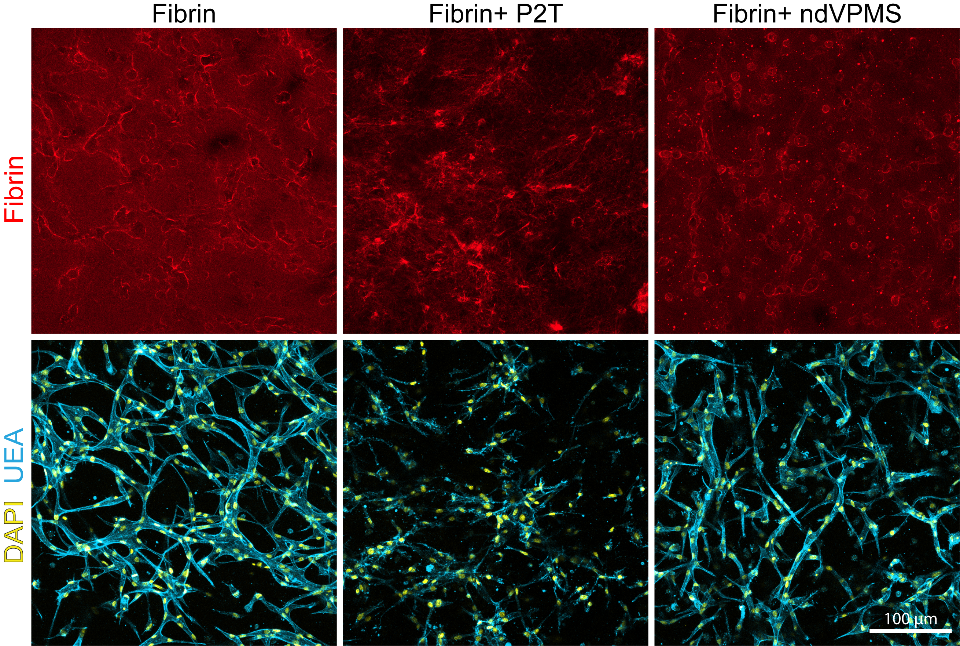

**Figure Supplement 1: P2T affects fibrin ultrastructures and vasculogenic assembly while ndVPMS does not.** Confocal fluorescent images of fibrin (top) and vascular networks (bottom) in fibrin gels spiked with P2T or ndVPMS.

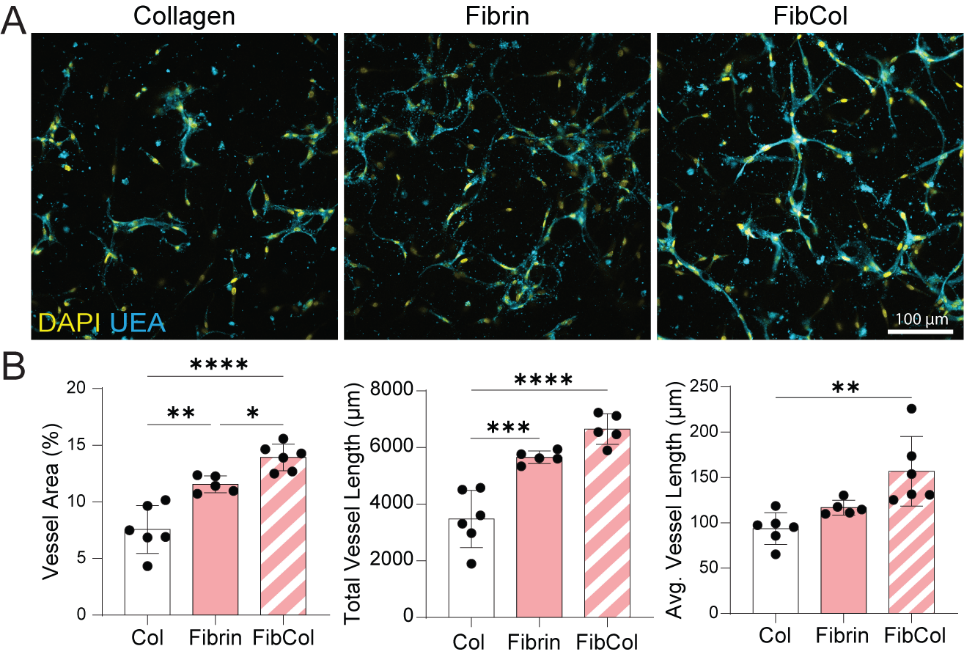

**Figure Supplement 2: FibCol best supports vascular network formation within GHCs.** (A) Confocal fluorescent images of vascular networks in GHCs with collagen, fibrin, and FibCol interstitial matrices. (B) Vessel area fraction, total vessel length, and average vessel length quantifications, respectively. *P < 0.05.

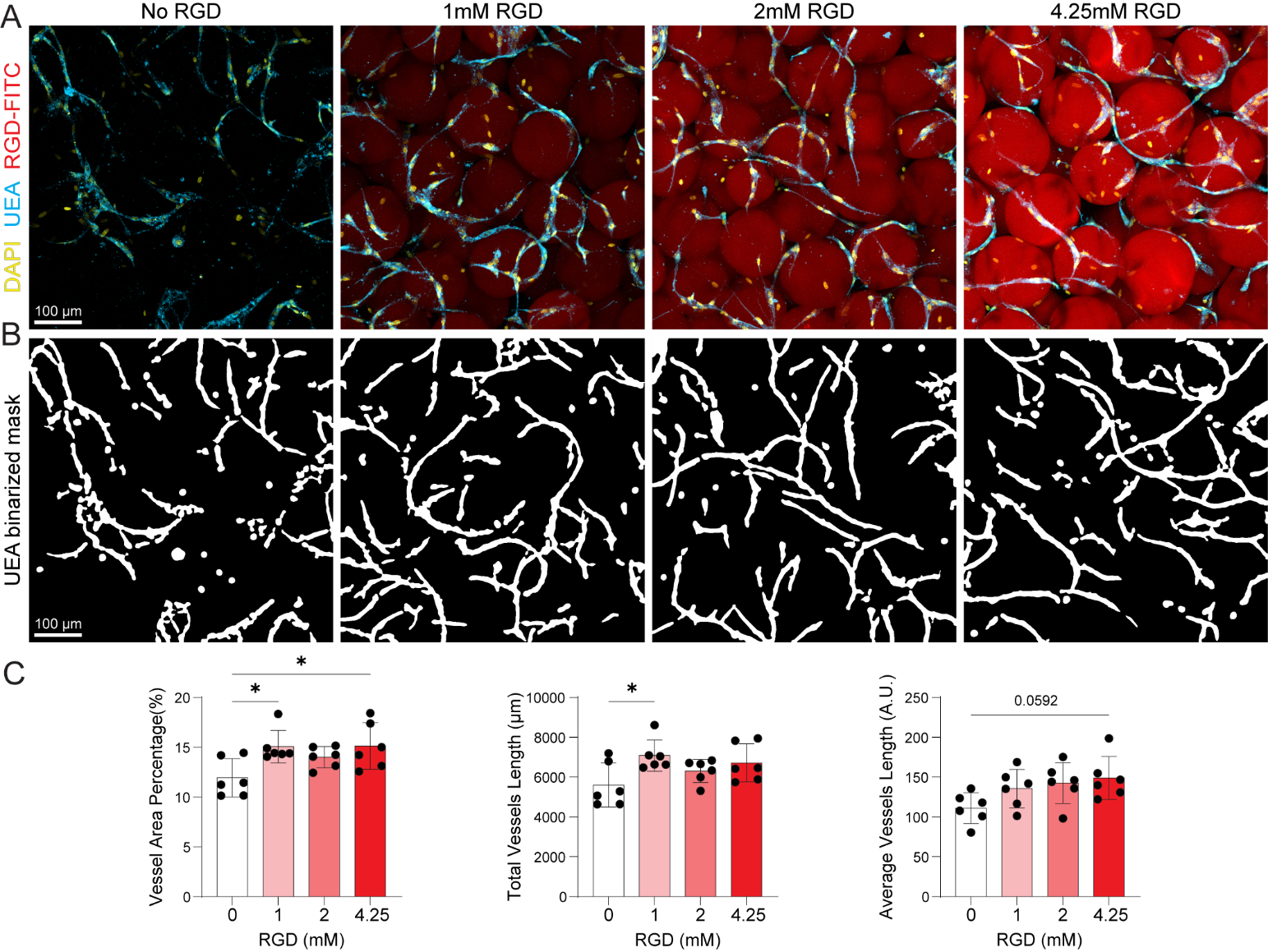

**Figure Supplement 3: RGD functionalization of microgels improves vascular network formation.** (A) Confocal fluorescent images of assembled vascular networks as a function of RGD treatment concentration. (B) Image analysis to identify and quantify vascular networks shown in (A). (C) Vessel area fraction, total vessel length, and average vessel length quantifications as a function of RGD treatment concentration. *P < 0.05.

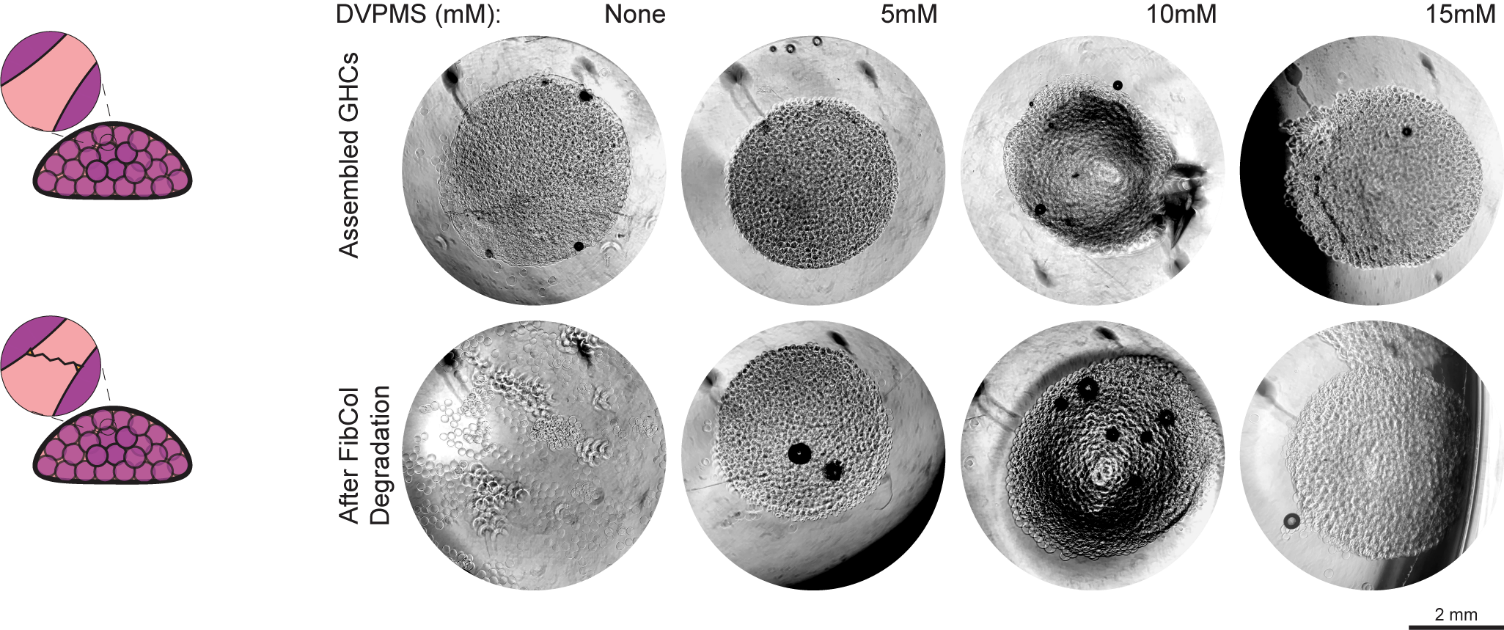

**Figure Supplement 4: Microgel interlinking is necessary for GHC scaffold integrity following collagen and fibrin matrix degradation.** (Left) Schematic overview of interstitial matrix degradation. (Right) Brightfield images of GHC scaffolds before and after interstitial matrix degradation as a function of interlinking agent DVPMS concentration.

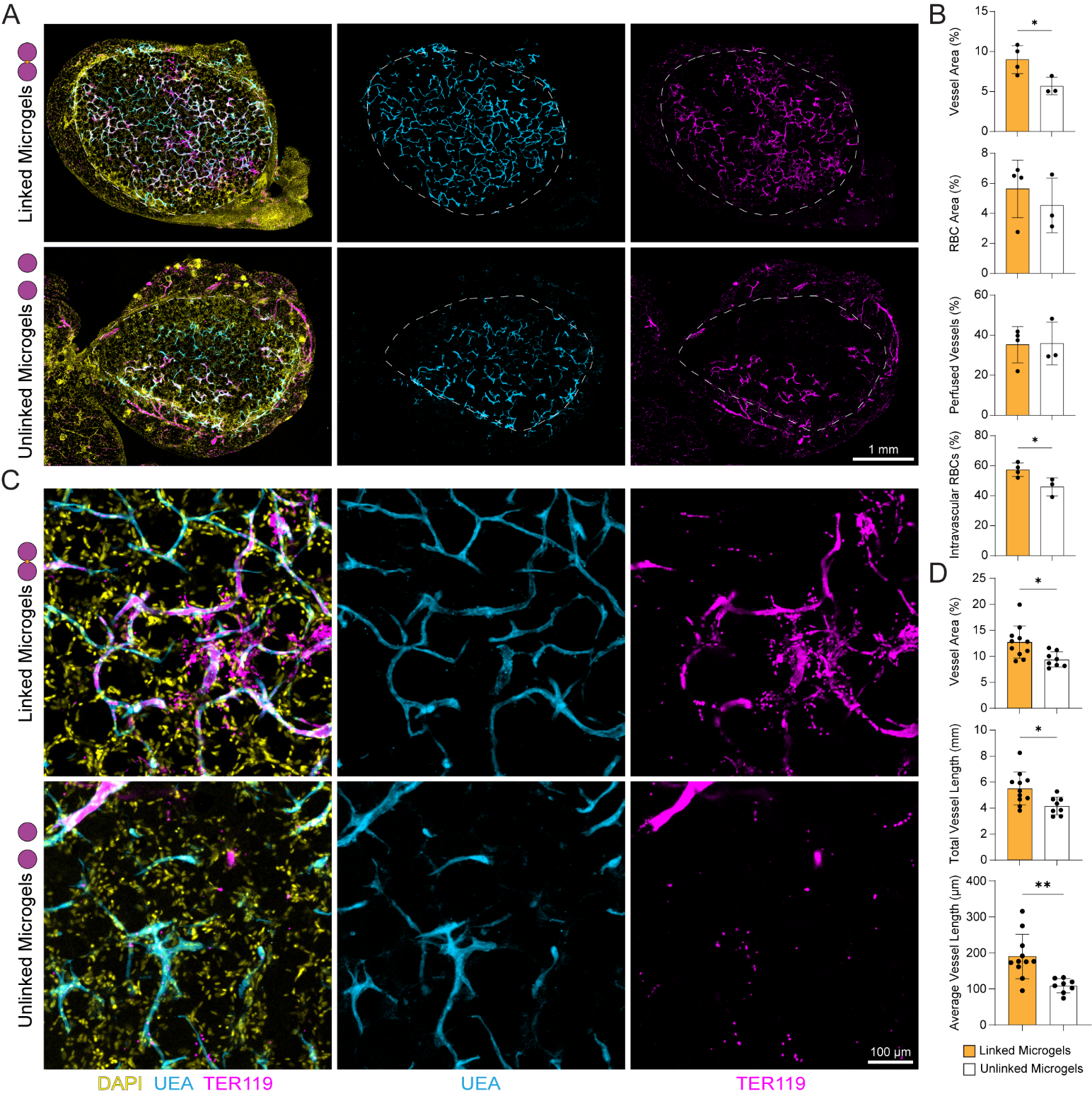

**Figure Supplement 5: Microgel interlinking improves the vascular integration of GHC grafts.** (A) Confocal max. intensity projections of interconnected and not connected GHC graft slices. (B) Graft vessel area fraction, RBC area fraction, perfused vessel fraction, and intervascular RBC fraction quantifications, respectively. (C) Fluorescent images of HUVEC networks in explanted grafts. (F) Vessel network area and total vessel length quantifications, respectively. *P < 0.05.
